# Harnessing Bilayer Biomaterial Delivery of FTY720 as an Immunotherapy to Accelerate Oral Wound Healing

**DOI:** 10.1101/2023.12.22.573096

**Authors:** Afra I. Toma, Daniel Shah, Daniela Roth, Jeremie Oliver Piña, Lauren Hymel, Thomas Turner, Archana Kamalakar, Ken Liu, Perry Bartsch, Leon Jacobs, Rena D’Souza, Dennis Liotta, Edward Botchwey, Nick J. Willett, Steven L. Goudy

**Affiliations:** Wallace H. Coulter Department of Biomedical Engineering, Georgia Institute of Technology, Emory University, Atlanta, GA, USA; Department of Pediatrics and Otolaryngology, Children’s Healthcare of Atlanta, Atlanta, GA, USA; Eunice Kennedy Shriver National Institute of Child Health and Human Development, National Institutes of Health, Bethesda, MD, USA; Department of Chemistry, Emory University, Atlanta, GA, USA; Phil and Penny Knight Campus for Accelerating Scientific Impact, University of Oregon, Eugene, OR, USA

## Abstract

Orofacial clefts are the most common craniofacial congenital anomaly. Following cleft palate repair, up to 60% of surgeries have wound healing complications leading to oronasal fistula (ONF), a persistent connection between the roof of the mouth and the nasal cavity. The current gold standard methods for ONF repair use human allograft tissues; however, these procedures have risks of graft infection and/or rejection, requiring surgical revisions. Immunoregenerative therapies present a novel alternative approach to harness the body’s immune response and enhance the wound healing environment. We utilized a repurposed FDA-approved immunomodulatory drug, FTY720, to reduce the egress of lymphocytes and induce immune cell fate switching toward pro-regenerative phenotypes. Here, we engineered a bilayer biomaterial system using Tegaderm™, a liquid-impermeable wound dressing, to secure and control the delivery of FTY720- nanofiber scaffolds (FTY720-NF). We optimized release kinetics of the bilayer FTY720-NF to sustain drug release for up to 7d with safe, efficacious transdermal absorption and tissue biodistribution. Through comprehensive immunophenotyping, our results illustrate a pseudotime pro-regenerative state transition in recruited hybrid immune cells to the wound site. Additional histological assessments established a significant difference in full thickness ONF closure in mice on Day 7 following treatment with bilayer FTY720-NF, compared to controls. These findings demonstrate the utility of immunomodulatory strategies for oral wound healing, better positing the field to develop more efficacious treatment options for pediatric patients.

**One Sentence Summary:** Local delivery of bilayer FTY720-nanofiber scaffolds in an ONF mouse model promotes complete wound closure through modulation of pro-regenerative immune and stromal cells.

## INTRODUCTION

Cleft lip and/or palate is one of the most common congenital anomalies, occurring in 1:940 live births(*1, 2*). Children typically require cleft lip surgery by three to six months, and palate surgery between six and twelve months of age(*3*). However, due to adverse healing following repair, up to 60% of patients have wound healing complications leading to oronasal fistula (ONF), an improper connection between the oral and nasal cavity(*4*). ONF can result in food lodgment, leading to bacterial accumulation and mucosal inflammation. High ONF occurrence is attributed to variability in surgical techniques, patient’s age and healing capability, as well as severity of the original injury. Future surgical attempts are only successful on average ∼50% of the time due to poor wound healing leading to infection, flap necrosis, hematoma formation, inadequate occlusion, and constant biomechanical tension(*5–7*). This can affect the child’s ability to eat, develop proper speech, and thus overall quality of life; hence, it is imperative to advance clinical therapeutics for the treatment of ONF.

The oral cavity presents a unique challenge in developing a biomaterial scaffold that can integrate to the host tissue environment due to the bacteria, saliva, and constant biomechanical forces. The current gold standard treatment in ONF repair involves using commercially available tissue grafts and can be categorized into two groups: allografts and autografts. Treatment with tissue grafts is beneficial when there is inadequate supply of local tissue for autografts, especially in large or severe oral mucosal wounds. However, use of these graft materials is an off-label treatment as they do not have FDA-approval specifically for mucosal replacement. Tissue grafts can augment oral wound healing to avoid ONF occurrence after palatoplasty but function solely as structural barriers and lack functional capabilities, often leading to graft failure(*8, 9*). Therefore, there is an unmet need to develop an implantable therapy which mitigates transplant rejection by leveraging oral wound healing cascades, reduces wound persistence, and can be FDA-approved for mucosal tissue repair.

FTY720 (Fingolimod) is an FDA-approved immunomodulatory drug currently prescribed as a once-daily oral drug treatment for relapsing multiple sclerosis (MS)(*10*). It acts as a Spingosine-1-Phosphate (S1P) receptor small molecule agonist that sequesters lymphocyte from secondary lymphoid organs to prevent an autoimmune response(*11*). S1P is a blood-borne bioactive sphingolipid abundant in endothelial cells, playing a key regulatory role in cell survival, trafficking, suppressing apoptosis, and nuclear signaling pathways(*12*). In vivo FTY720 is synthesized into its bioactive form through sphingosine kinase-2 (SPHK2), located in the cell nucleus, and has a higher affinity than S1P to bind extracellularly with one of five known G-protein-coupled receptors termed S1PR_1-5_(*13*). Upon binding, FTY720-Phosphate (FTY720P) internalizes the receptor, activating intracellular targets and simultaneously induces degradation of S1PR, preventing lymphocyte egression and driving immune cell fate switching towards a pro-regenerative state (*11*). To use FTY720 as an immunoregenerative candidate for ONF treatment, we previously fabricated a nanofiber (NF) scaffold loaded with FTY720, serving as a 3D template to direct oral tissue regeneration as a local, transmucosal drug delivery system. Preliminary results from our lab reported that FTY720-nanofiber scaffolds (FTY720-NF) selectively recruited pro-regenerative monocytes and altered gene expression of pro-regenerative cytokines and transcription factors(*14*). While promising, we recognize crucial limitations within our previous research of FTY720-NF as a clinically translatable immunotherapy for ONF healing.

In this work, we continued our development of a novel, pro-regenerative immunotherapeutic strategy for oral wound healing by engineering a bilayer FTY720-NF system to locally modulate the oral immune response following cleft palate repair to reduce post-operative ONF occurrence. There remained key technological innovations that are now addressed in the present study to improve scaling and translation of the technology toward clinical application. To scale the approach and ensure consistent, sustained FTY720 delivery, we 1) evaluated drug dose response, release kinetics, and biodistribution of FTY720 both in vitro and in vivo, 2) developed novel material strategies using a Tegaderm™ (TD) bilayer adhesive system to enhance retention of the scaffold in the local ONF site and 3) examined the function of pro-regenerative immune cells that contribute to successful tissue closure and ONF healing. Herein, we demonstrated how FTY720-NF acts as a novel form of pro-regenerative immunotherapy for oral wound healing, paving the way toward safe and efficacious treatment options for patients affected by cleft palate who often experience post-operative oral wound healing complications.

## RESULTS

### FTY720-NF sustains drug release over 7 days

In order to design an efficacious FTY720 delivery system that will allow sustained released for up to 7 days, we fabricated electrospun nanofibers using poly(lactide-co-glycolide acid (PLGA) and polycaprolactone (PCL) and analyzed FTY720-NF release kinetics and drug loading capacity in vitro. We characterized FTY720-NF using high-performance liquid chromatography-tandem mass spectrometry (LC/MS) to confirm drug release kinetics and loading efficiency in scaffolds loaded with 0mg (-CTRL), 1:120, and 1:60 drug:polymer concentration ratio (*Fig. 1C-E*). Our data revealed that 1:60 ratio cumulatively released 1.37µg ± 0.070µg FTY720 per 1.5mm diameter NF with 85.32% loading efficiency over 7d time course. In 1:120 ratio, 0.27µg ± 0.033µg FTY720 per NF was released with 75.23% loading efficiency over 48hr time course. All future characterization and in vivo studies comprised of the 1:60 concentration as it significantly enhanced drug release compared to the 1:120 ratio (*P*=0.0048). Ordinary one-way ANOVA with Tukey’s multiple comparisons was conducted for statistical analysis and data is presented as mean ± S.E.M with significance at *P*<0.05. n = 3–5 scaffolds per drug loading group.

**Fig. 1.**
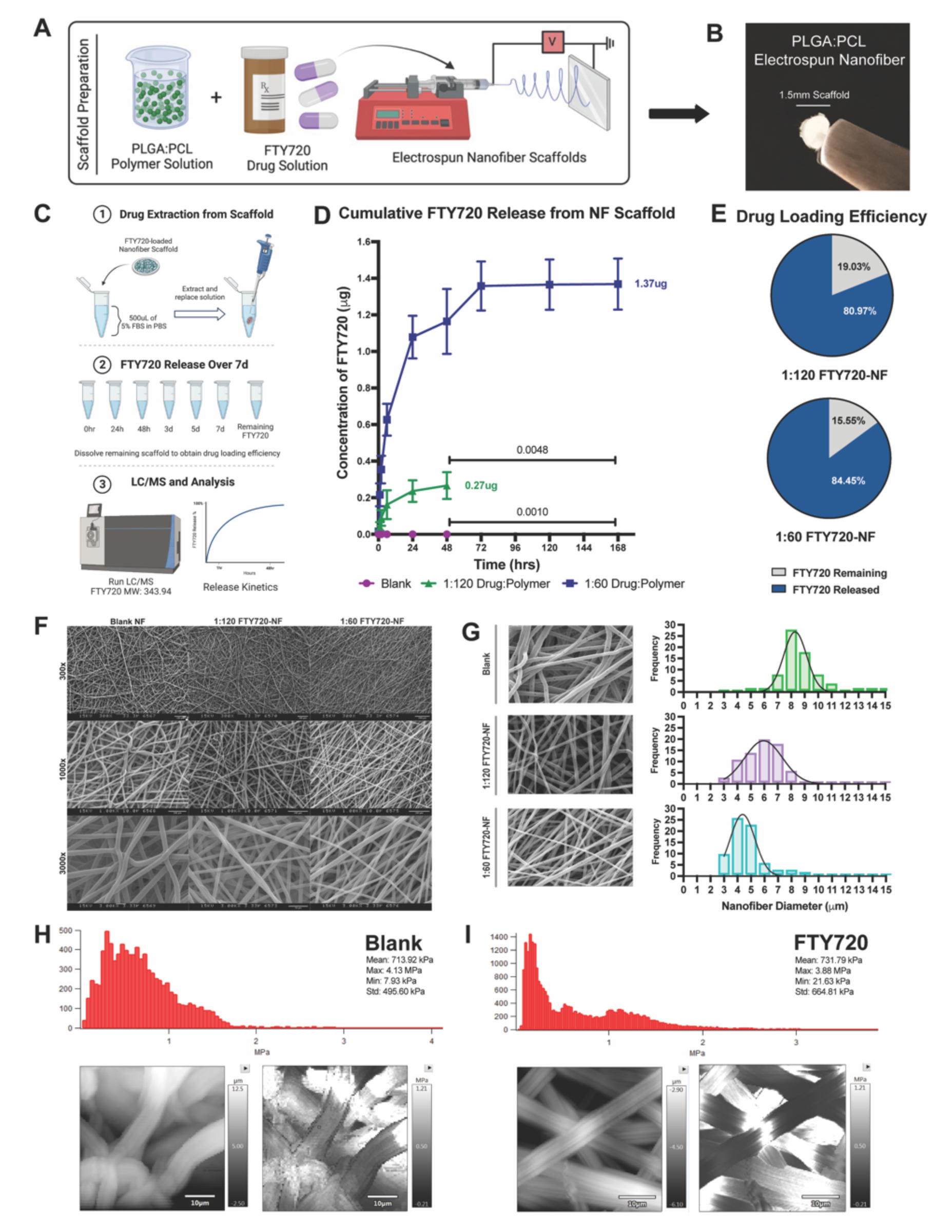
FTY720-NF sustained drug release for 7d and scaffolds retain similar mechanical properties between unloaded and drug-loaded nanofibers. **A-B**) FTY720 drug solution was coupled with PLGA:PCL polymers, electrospun into nanofiber, and biopsy-punched into 1.5mm diameter scaffolds. **C-E**) LC/MS reveals cumulative drug release and total drug loading efficiency from blank, 1:120, and 1:60 FT720:polymer weight scaffolds over 48hr or 7d. 1:60 drug scaffolds cumulatively released 1.37ug FTY720 with 84.45% release efficiency. **F-G**) SEM assessed morphology of blank, 1:120, and 1:60 FT720-NF scaffolds. **H-I**) AFM confirms that stiffness of blank and 1:60 FTY720-NF was equivalent regardless of incorporation of drug.

We assessed morphology and stiffness between unloaded and loaded scaffolds using scanning electron microscopy (SEM) and atomic force microscopy (AFM) to ensure fabrication consistency and reproducibility (*Fig. 1F-I*). Scanning electron micrographs revealed a similar randomly oriented morphology in all scaffold groups with nanofiber width of 8.69µm ± 0.19µm for blank, 5.75µm ± 0.15µm for 1:120 FTY720-NF, and 4.65µm ± 0.15µm 1:60 FTY720-NF. We performed AFM to evaluate the surface of the nanofibers with high resolution and confirmed that the topography of 1:60 FTY720-NF was quantitatively consistent with unloaded drug scaffold. Young’s modulus values were obtained over multiple contact points on the nanofibers, showing similar distribution profiles between unloaded and loaded scaffolds. For AFM surface topology and stiffness maps, lighter regions represent areas of greater height and stiffness for blank and FTY720-NF. Histograms of Young’s moduli (MPa) were obtained over multiple contact points on unloaded and 1:60 FTY720-NF. Results showed that blank scaffolds produced Young’s Moduli of 713.92kPa ± 495.60kPa and max at 4.13MPa. Similarly, FTY720-NF had a Young’s Moduli of 731.79kPa ± 664.81kPa and max at 3.88MPa. These techniques established that the fabrication process is highly reproducible and there are no significant effects of the incorporation of the drug on scaffold morphology and topology.

### Bilayer adhesive scaffold for FTY720 delivery accelerates ONF wound closure by D7

We hypothesized that fixation of FTY720-NF to the hard palate mucosa would improve transdermal drug delivery and allow FTY720 to locally modulate S1P receptors following scaffold implantation in our previously established in vivo ONF surgery model (*Fig. 2A-B*)(*14*). Various scaffold adhesive methods were compared to secure FTY720-NF to site of ONF injury on the palate, and adhesives were assessed for duration of adherence to oral palate (*Supplemental Fig. 1A*). 3M Vetbond™ and Oasis LiquiVet 8-second Rapid Tissue Adhesive helped adhere the scaffold directly to the oral tissue for >3d following implantation. However, an LC/MS study revealed no FTY720 release in vitro when scaffold is directly adhered with tissue glue, potentially due to the glue seeping into the nanofibers and preventing drug diffusion (*Supplemental Fig. 1B*). Tissue adherence with TBM Corporation Ora-Aid, GlaxoSmithKline Poligrip® Denture Adhesive, and 8P-T Mucosal Hydrogel caused the FTY720-NF to dislodge from the palate within 2 hours, making it inadequate to allow sufficient drug release from the scaffold to the local tissue. 3M Nexcare Tegaderm™ (TD) is a transparent polyurethane film that is ubiquitously used in the clinical setting to protect I.V. catheter sites and wounds. TD is commercially available as a sterile, semi-impermeable thin film that is occlusive to liquid, bacteria, and small molecules yet can easily exchange water vapor, oxygen, and carbon dioxide. We layered TD above the FTY720-NF implanted at the ONF site and thinly coated the TD with sterile tissue glue, securing the scaffold to the oral mucosa (*Fig. 2C-D*). This bilayer adhesive system provided the longest duration of scaffold attachment for at least 8h and up to 24h, facilitating localized drug release to the oral mucosa (*Fig. 2E*). TD bilayer acted as a physical barrier to adhere FTY720-NF to the oral mucosa, providing a waterproof film while maintaining a moist environment for wound healing.

**Fig. 2.**
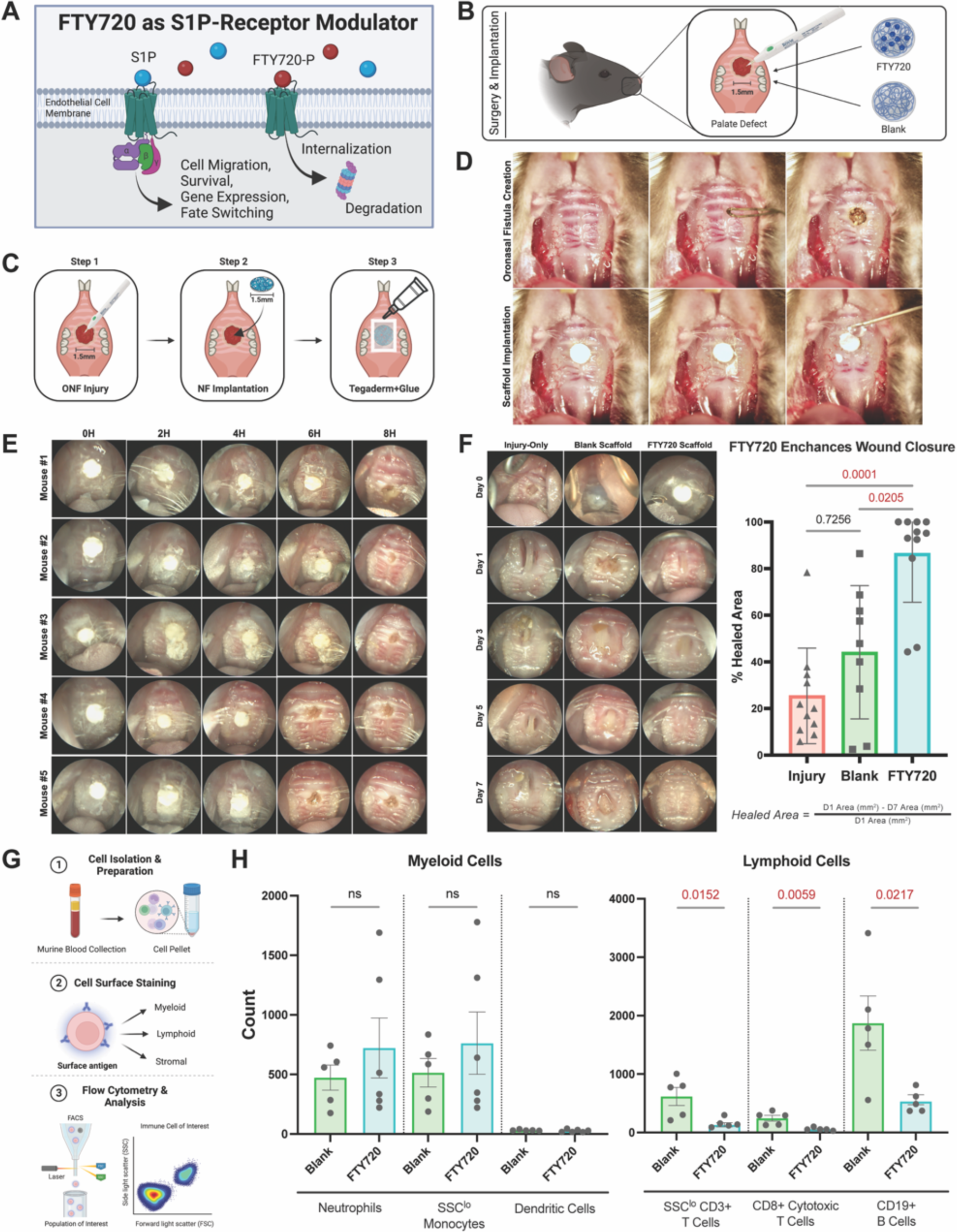
Tegaderm™ bilayer adhesive system enhanced retention of FTY720-NF in the ONF site and enhanced oral tissue regeneration by D7. **A)** Mechanism of action for FTY720 as an S1P receptor modulator by inducing receptor internalization and degradation, preventing receptor-mediated cell migration. **B-D)** FTY720 or control nanofiber scaffolds were implanted at the site of ONF injury in C57BL/6 mice (n=9-11) on D0 and secured with a bilayer adhesive system using Tegaderm™. **E)** Scaffold adhesive to oral tissue using Tegaderm™ was recorded up 24hr post-implantation. **F)** Endoscope images of wound closure were taken at D1, 3, 5, and 7, and injury size was measured using Image J software. Total wound closure was compared between injury-only, blank, and FTY720-treated mice. Mice treated with FTY720-NF showed significant wound closure compared to controls (*P*=0.0001). **G-H)** Functional testing of FTY720-NF resulted in acute lymphopenia of blood lymphoid cells at 24h, indicating that FTY720 was successfully delivered via scaffold implantation.

Bilayer scaffolds, either blank or with FTY720, were implanted at the site of ONF injury in mice on Day 0. Injury-only mice were used as negative control to establish a baseline critically sized defect with persistent ONF at D7. Endoscopic images of wound closure were taken at D1, 3, 5, and 7, and injury size was measured using ImageJ software (*Fig. 2F*). Endoscopic measurements of wound size showed significantly improved wound closure in mice treated with FTY720-NF by D7 when compared to control groups (blank, *P*=0.0205 and injured mice, *P*=0.0001). As expected, there was no significant difference between blank and injury-only mice (*P*=0.7256) confirming that NF alone does not promote significant ONF closure without FTY720. These results demonstrated that mice treated with FTY720-NF exhibit accelerated healing with full ONF closure by D7. Kruskal-Wallis one-way ANOVA was conducted for statistical analysis and data is presented as mean wound size ± S.E.M with significance at *P*<0.05. n = 9-11 C57BL/6 mice (age 4-6wks) per treatment group.

### Mice treated with FTY720-NF exhibit acute lymphopenia at 24H

FTY720 acts primarily on lymphocytes with reversible transient lymphopenia(*15*). To evaluate the systemic impact of FTY720 on circulating lymphocytes, we collected blood from mice at 24h as a terminal timepoint and count was compared using flow cytometric techniques (*Fig. 2G-H*). Each blood sample was analyzed for CD45^+^ leukocytes and gated for innate and adaptive immune cell subsets. Significant differences were evaluated using a two-tail, unpaired t-test and is presented as mean counts ± S.E.M with significance at *P*<0.05. n= 5-6 C57BL/6 mice were used per treatment group.

Within CD11b^+^ myeloid cell population, analysis revealed no significant difference in blood cell counts between treatment groups for Ly6G^+^ neutrophils, MerTK^−^ monocytes, and CD11c^+^ dendritic cells. These results indicate that blood myeloid cells were not affected by FTY720-NF at 24h post treatment and suggest that FTY720 may influence myeloid cell functions through fate switching, cell survival, or cytokine production. In contrast, when gating for CD11b^−^ lymphoid immune cells, mice treated with FTY720-NF exhibited significantly lower counts of circulating CD3^+^ T-cells (*P*=0.0152), CD8^+^ cytotoxic T-cells (*P*=0.0059) and CD19^+^ B-cells (*P*=0.0217) compared to control mice at 24h. Flow cytometric data confirmed acute lymphopenia in FTY720-NF mice at 24h, concluding that the scaffold-adhesive system allowed for successful transmucosal drug absorption at the local implantation site and effectively decreased ingress of lymphocytes into the blood.

### Treatment with FTY720-NF causes pseudotime shift in phenotypic trajectories of myeloid immune cells during ONF healing

During the early stages of ONF wound healing, we evaluated the ingress of immune cells and identified pro-regenerative cell subtypes recruited to the wound bed following FTY720-NF treatment (*Fig. 3A*). Following FTY720-NF or control bilayer scaffold implantation, hard palate tissue was collected across all timepoints (Days 1, 3, 5, and 7) to identify local immune cells at the injury site. Palatal tissue explanted from naïve and injury-only mice served as control samples. Flow cytometry and multi-dimensional immunophenotyping analyses quantified immune cell count across three immune cell classes: myeloid, lymphoid, and stromal cells (*Fig. 3-5*). All flow cytometric studies and subsequent analyses were conducted with n=11-12 biologically independent C57BL/6J mice per experimental group, as outlined in *Material and Methods*.

**Fig. 3.**
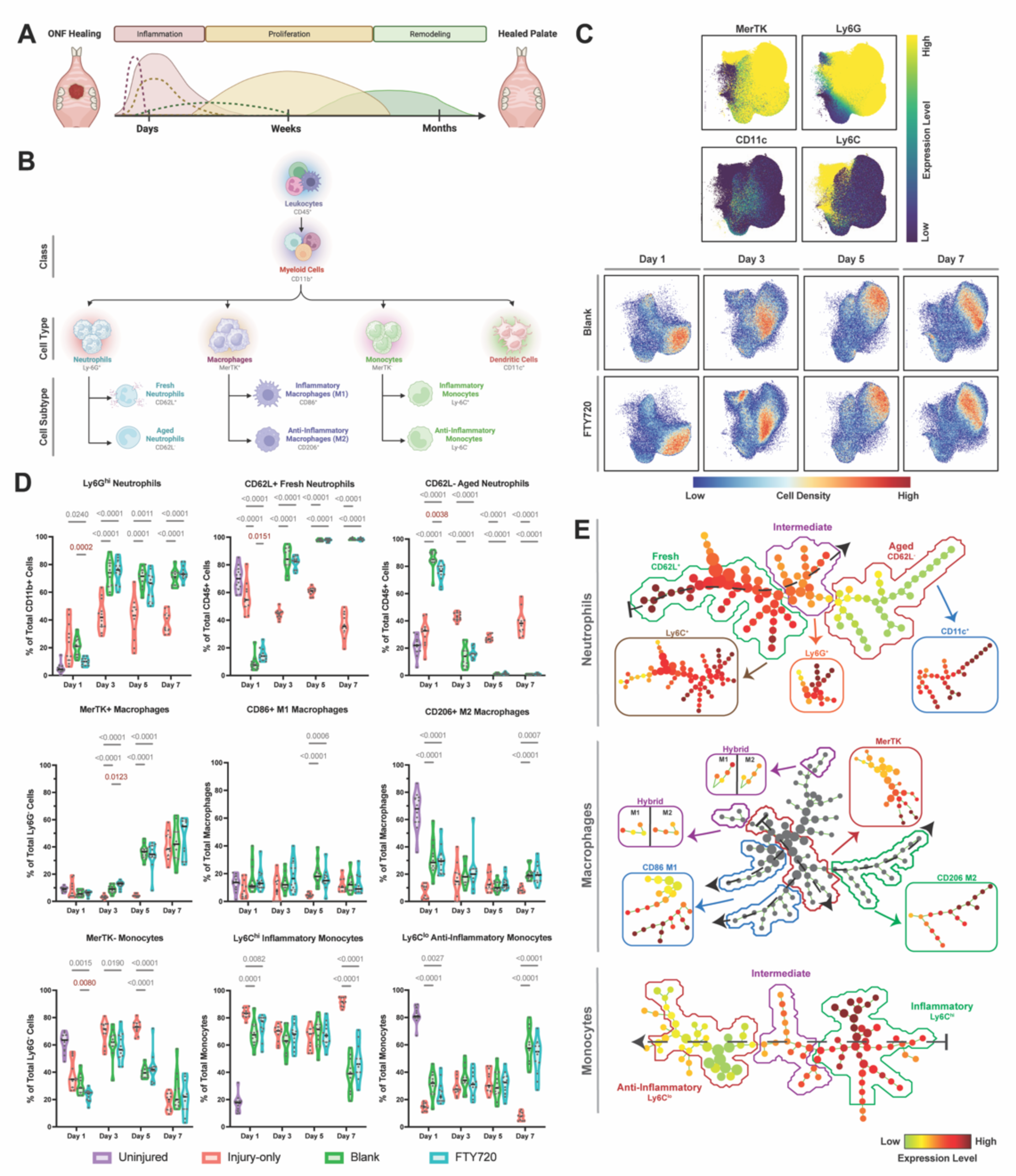
Treatment with FTY720-NF reveals myeloid immune cell recruitment similar to healing cascade and a pseudotime trajectory toward pro-regenerative phenotypes. **A)** Schematic of spatiotemporal phases of oral wound healing: inflammation, proliferation, and remodeling. **B)** Gating strategy used within flow cytometric studies to identify neutrophils, macrophages, and monocytes cell types deriving from myeloid cell class. **C)** UMAPs provide a visual representation of single cells by their myeloid cell expression, ranging from dark violet (low expression) to yellow (high expression) and myeloid cell density following blank or FTY720-NF treatment on Days 1, 3, 5, and 7, ranging from blue (low expression) to red (high expression). **D)** Flow cytometric data reveals significantly lower concentration of neutrophils (*P*<0.0002) and monocytes (*P*<0.0080) following FTY720-NF treatment on D1 but significantly higher macrophage percentage (*P*<0.0123) by D3 when compared to blank scaffolds. **E)** Neutrophil SPADE dendrogram depicts transition of cells from fresh trajectory to intermediate and aged phenotypes. Macrophage and Monocyte SPADE dendrograms reveal that inflammatory trajectory transition to pro-regenerative phenotypes by D7. Significance was evaluated in a 2-way ANOVA with Tukey’s multiple comparison with *P*<0.05 for n=11-12 biologically independent C57BL/6J mice per experimental group.

The flow panel was designed for myeloid cell class and subtypes identified by CD45^+^CD11b^+^ antibodies, a canonical myeloid cell surface leukocyte marker used to delineate myeloid-derived immune cell types (*Fig. 3B*). Ly6G is commonly used as a marker for neutrophils and can be further differentiated using CD62L for fresh vs aged neutrophils(*16*). Macrophages were identified as CD64^+^MerTK^+^, a receptor tyrosine kinase, and polarization state was characterized further using markers CD86^+^ for pro-inflammatory M1 polarization or CD206^+^ for alternatively activated M2 macrophages(*17*). The MerTK^−^Ly6C^+^ antigen differentiates between classical Ly6C^hi^ or non-classical Ly6C^lo^ monocytes present in murine palatal tissue(*18*).

UMAPs were constructed with OMIQ.ai using flow cytometry data from all CD11b^+^ myeloid cells from blank or FTY720-NF treated mice and aggregated across all timepoints (*Fig. 3C*). Plots provide a visual representation of single cells by their surface marker profiles, with proximity between cells reflecting cellular phenotypes. Surface marker expression values among myeloid cell types are represented on the UMAP, ranging from dark violet (low expression) to yellow (high expression). A CD11b^+^ myeloid cellular event is represented as a dot on the UMAP with distinct cell phenotypes between mice treated with FTY720-NF compared to control, ranging from blue (low cell density) to red (high cell density). On D1, there was a qualitative shift in Ly6G^+^ neutrophil expression with higher density in the blank group compared to FTY720. Additionally, on D3, there was higher MerTK^+^ expression in FTY720-NF compared to control, indicating that treatment with FTY720 can modulate myeloid cell recruitment and expression during early stages of healing.

Myeloid immune cells were gated, and cell counts were analyzed across all timepoints for mice (n=11-12) in treatment groups: injury-only, blank, and FTY720 (*Fig. 3D*). Immune cell types are shown as percentage of parent population with mean ± S.E.M. 2-way ANOVA with Tukey’s multiple comparison revealed significance at *P*<0.05 between treatment groups for neutrophil, macrophage, and monocyte immune cell types. On D1, mice treated with FTY720-NF presented significantly lower percentage of all neutrophils compared to control mice (blank, *P*=0.0002 and injury-only *P*=0.0240). This confirms that neutrophils extravasate early in the healing cascade as part of innate immunity involved in pathogen debridement which contributes to the activation of macrophages to resolve inflammation(*19, 20*). However, FTY720-NF mice had significantly higher CD62L^+^ fresh neutrophil percentages compared to blank (*P*=0.0151) but had lower CD62L^−^ aged neutrophil recruitment (*P*=0.0038) on D1. Following an injury, freshly released neutrophils migrate from bone marrow into the tissue and undergo changes in their phenotype and morphology every four hours, whereas, neutrophils that have aged in the circulation are eliminated at the end of the resting period via phagocytic macrophages(*21, 22*). Mice treated with FTY720-NF produced a significantly higher percentage of macrophages on D3 when compared to controls (blank, *P*=0.0123 and injury-only, *P*=0.0001), suggesting that FTY720-NF further induces macrophage functions of antigen presentation, phagocytosis, and regulation of chemokine and cytokine secretion. However, no significant difference between groups was indicated in the M1 and M2 macrophage polarization group across all timepoints. Mice that received FTY720-NF presented with a significantly lower percentage of monocytes on D1 compared to control mice (blank, *P*= 0.0080 and injury-only, *P*=0.0015). Monocyte subsets did not reveal any significant difference between treatment groups, indicating that FTY720 primarily acted on macrophages following monocyte differentiation in the tissue.

We generated a SPADE dendrogram from all CD11b^+^ myeloid cells isolated from palate tissue of mice treated with blank or FTY720-NF across all timepoints (*Fig. 3E*). The neutrophil SPADE dendrogram is color annotated into three subpopulations of Ly6G^+^ neutrophils containing CD62L^+^ fresh, intermediate, and CD62L^−^ aged expression. From left to right, pseudotime projection of neutrophil SPADE demonstrated a phenotypic transition from fresh neutrophils at D1 to intermediate and aged expression by D7 following ONF injury. Next, the macrophage SPADE dendrogram delineated three distinct subpopulations of macrophages containing M1 inflammatory, hybrid, and M2 anti-inflammatory expression. Moving down the dendrogram, as indicated by dotted black arrows, the clustered nodes split into 2 separate branches, M1 trajectory on the left and M2 trajectory on the right with hybrid macrophage trajectory in intermediate nodes. Lastly, the monocyte SPADE dendrogram clustered into 3 marked trajectories of Ly6C^lo^ anti-inflammatory, intermediate Ly6C, and inflammatory Ly6C^hi^ nodes. Most monocytes present in the palate at D1 were clustered in nodes with inflammatory trajectory and transitioned to anti-inflammatory trajectory by D7. Our data shows that myeloid cells with inflammatory trajectories early in the wound healing cascade transition to anti-inflammatory phenotypes by D7, further demonstrating that FTY720-NF can promote a pro-regenerative healing environment following ONF injury. These findings are consistent with successful wound healing cascades indicating a shift in immune cell state transition toward a pro-regenerative phenotype. S1P modulates macrophage responses according to the local environment so that it not only promotes the production of M2, but also of M1-associated macrophage markers.

### FTY720-NF induces lymphopenia at ONF injury with distinct activation of T cell subsets to regulate lymphoid cell phenotype

Evaluation of palatal tissue for lymphocyte recruitment was performed by harvesting tissues as described above and using a flow panel designed for the lymphoid cell class identified by CD45^+^CD11b^−^ antibodies (*Fig. 4B*). CD3 is a T cell receptor complex generated by thymocytes that can be further differentiated into CD4^+^ T helper (T_h_) and CD8^+^ T cytotoxic cells (T_c_) (*23*). Antibodies against adhesion molecule CD44, a hyaluronan receptor, delineates activated T_h_ cells into subpopulations of CD44^−^CD62L^+^ naïve (T_N_), central memory (T_CM_), and CD44^+^CD62L^−^ effector memory (T_EM_) cells(*24*). B cells are the precursors of antibody-secreting cells and are identified by CD19^+^ found on the cell surface(*25*). CD161^+^, a C-type lectin-like receptor that is one of the earliest markers expressed in natural killer (NK) cell development. Lastly, CD11c^+^ is a marker for classical dendritic cells (DCs) and can be expressed under CD11b^+^ myeloid and CD11b^−^ lymphoid lineage(*26, 27*).

**Fig. 4.**
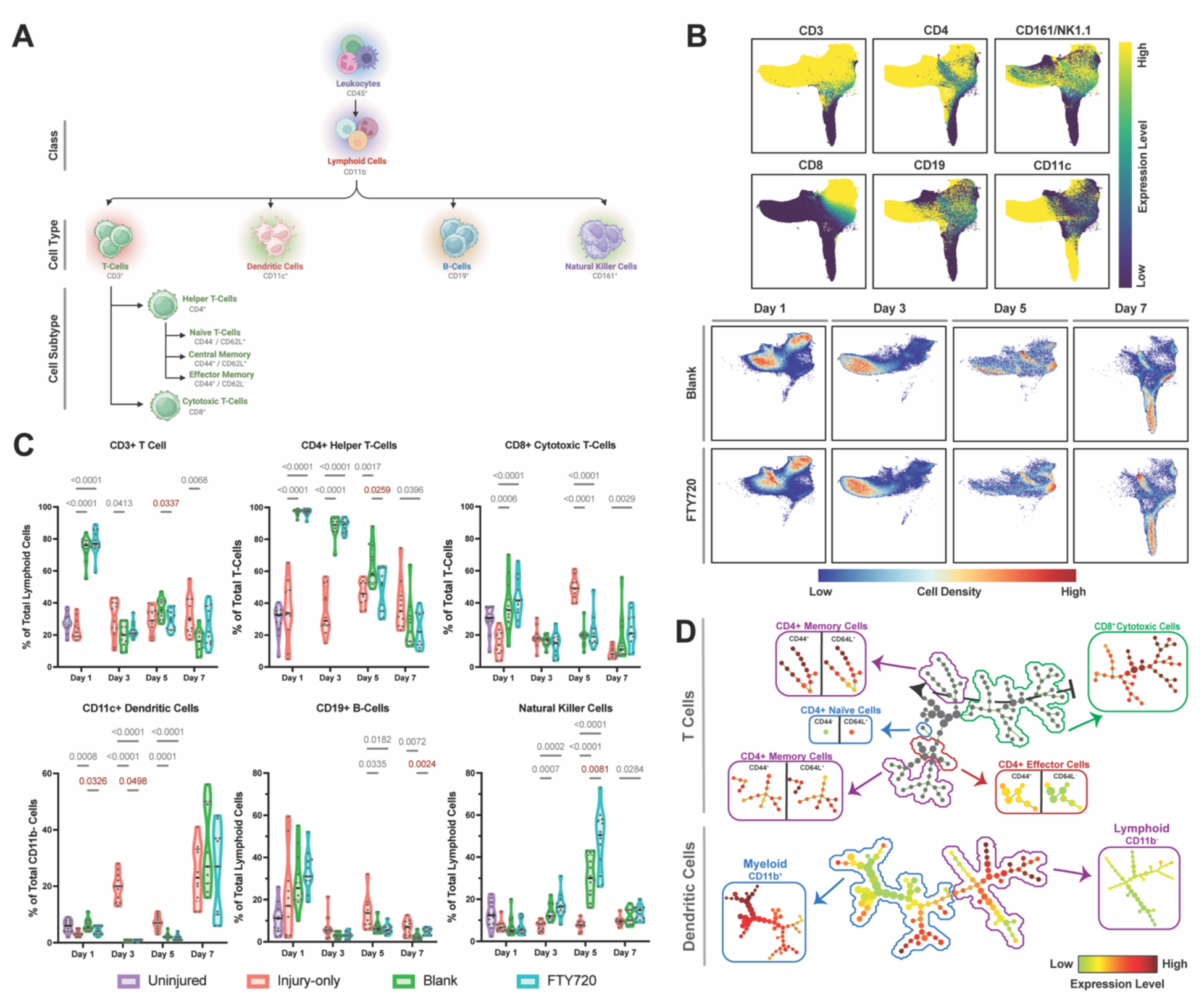
Treatment with FTY720-NF reveals lymphopenia in T cells with increased antibody-producing B cells and increased activation of lymphoid-derived DCs. **A)** Gating strategy used within flow cytometric studies to identify T cells, DCs, B cells, and natural killer cells deriving from lymphoid cell class. **B)** UMAPs provide a visual representation of single cells by their lymphoid cell expression, ranging from dark violet (low expression) to yellow (high expression) and lymphoid cell density following blank or FTY720-NF treatment on Days 1, 3, 5, and 7, ranging from blue (low expression) to red (high expression). **C)** Flow cytometric data reveals significantly lower concentration of DCs on D1 (*P*<0.0326) and D3 (*P*<0.0498) following FTY720-NF treatment. By D5, significantly lower concentrations were recorded for CD3^+^ T cells (*P*<0.0337), CD4^+^ T_h_ cells (*P*<0.0259), and NK cells (*P*<0.0081) compared to control scaffold. Higher concentration of CD19^+^ B cells (*P*<0.0024) were recorded by D7 following FTY720-NF treatment. **D**) T cell SPADE dendrogram shows the transition from T_c_ to T_CM_ expression. DC dendrogram reveals higher activation of lymphoid-derived DCs following ONF injury. Significance was evaluated in a 2-way ANOVA with Tukey’s multiple comparison with *P*<0.05 for n=11-12 biologically independent C57BL/6J mice per experimental group.

UMAPs were constructed using OMIQ.ai of all CD11b^−^ lymphoid cells from blank and FTY720-NF treated mice across all timepoints. Surface marker expression values among lymphoid cell types are represented on the UMAP ranging from dark violet (low expression) to yellow (high expression) (*Fig. 4C*). Cell density of lymphoid cells between blank and FTY720-NF groups is depicted in blue (low cell density) to red (high cell density). On Days 1 and 3, there is a higher cell density of CD3^+^ T cells, T_h_, and T_c_ cells compared to other days, with specifically higher density in the blank group compared to FTY720-NF. In the days following, there is higher expression of NK cells and DCs in both treatment groups.

Lymphocyte counts from flow cytometric data were analyzed across all timepoints for C57BL/6J mice (n=11-12) in all treatment groups: injury-only, blank, and FTY720-NF. Cell types were analyzed as a percentage of the parent population with mean counts ± S.E.M. 2-way ANOVA with Tukey’s multiple comparison revealed significant difference at *P*<0.05 between treatment groups for T cells, B cells, NK cells, and DCs (*Fig. 4D*). On Days 1 and 3, FTY720-NF treated groups exhibited a significantly lower percentage of DCs compared to control mice (blank, *P*=0.0326 and injury-only, *P*=0.008). Similarly, at D5 treatment of mice with FTY720-NF exhibited acute lymphopenia with significantly lower CD3^+^ T cells (*P*=0.0377) and CD4^+^ T_h_ cells (*P*=0.0259) with no significant difference to T_c_ cell percentage. By D5, a significantly higher percentage of NK cells were measured following FTY720-NF treatment compared to control mice (blank, *P*=0.0081 and injury-only, *P*=0.0001). By D7, FTY720-NF treatment produced a higher percentage of CD19^+^ B-cells (*P*=0.0024) compared to controls. Our data supports that DC presence on D1 likely control immune response by activating lymphocytes to tolerize T cells to self-antigens that are innate to the body, thereby minimizing autoimmune reactions in inflamed settings(*27, 28*). It is known that T cells fate switch to a state favoring egress over retention by upregulating S1PR_1_, so our data validates that a decrease in T cell percentage following treatment with FTY720 is due to T cells dependence on S1PR_1_-mediated egress(*29, 30*). A significant decrease in NK cells percentages following FTY720-NF probably limited their cytotoxic and cytokine-producing effector functions on the immune responses of DCs, macrophages, T cells and endothelial cells(*31*).

A SPADE dendrogram was generated from all CD11b^−^ lymphoid cells isolated from mice treated with blank or FTY720-NF (*Fig. 4E*). The T cell SPADE dendrogram is color depicted into four subpopulations of CD3^+^ cells containing T_c_ and T_h_ with distinction in T_N_, T_CM_, and T_EM_ cell phenotypes. From right to left, pseudotime projection of SPADE revealed a phenotypic transition from T_c_ cells at D1 to T_CM_ by D7 following ONF injury. However, the dendrogram revealed two distinct populations of T_CM_ cells across all time points. The dendritic cells SPADE dendrogram revealed two clusters of CD11c^+^ cells, denoting CD11b^−^lymphoid and CD11b^+^ myeloid lineage. SPADE showed higher expression of lymphoid-derived DCs following ONF injury when compared to DCs with myeloid lineage. These data suggest that during ONF healing, there is greater contribution from CD11b^−^ DCs that coincides with FTY720-mediated oral tissue closure.

### FTY720-NF regulates excessive stromal cell recruitment compared to injury-only mice

During wound repair, the extracellular matrix (ECM) is reconstituted by stromal cells which are non-immune cells involved in the repair of the cell-derived matrix through production of granulation tissue/stroma, vascularization, and differentiation of fibroblasts. To investigate stromal cell recruitment following FTY720-NF treatment, stromal cell subtypes were identified by CD45^−^ antibodies (*Fig. 5B*). CD31 is a surface marker for endothelial cells, which are hallmark cells facilitating angiogenesis and neovascularization during the proliferative phase. As new blood vessels and granulation tissue are laid down, CD326^+^ keratinocytes further drive cell proliferation and migration. Fibroblasts, identified by fibroblast activation protein alpha (FAPα), construct new ECM to support cell ingrowth, blood vessels to supply oxygen, and nutrients to sustain cell metabolism(*32*).

**Fig. 5.**
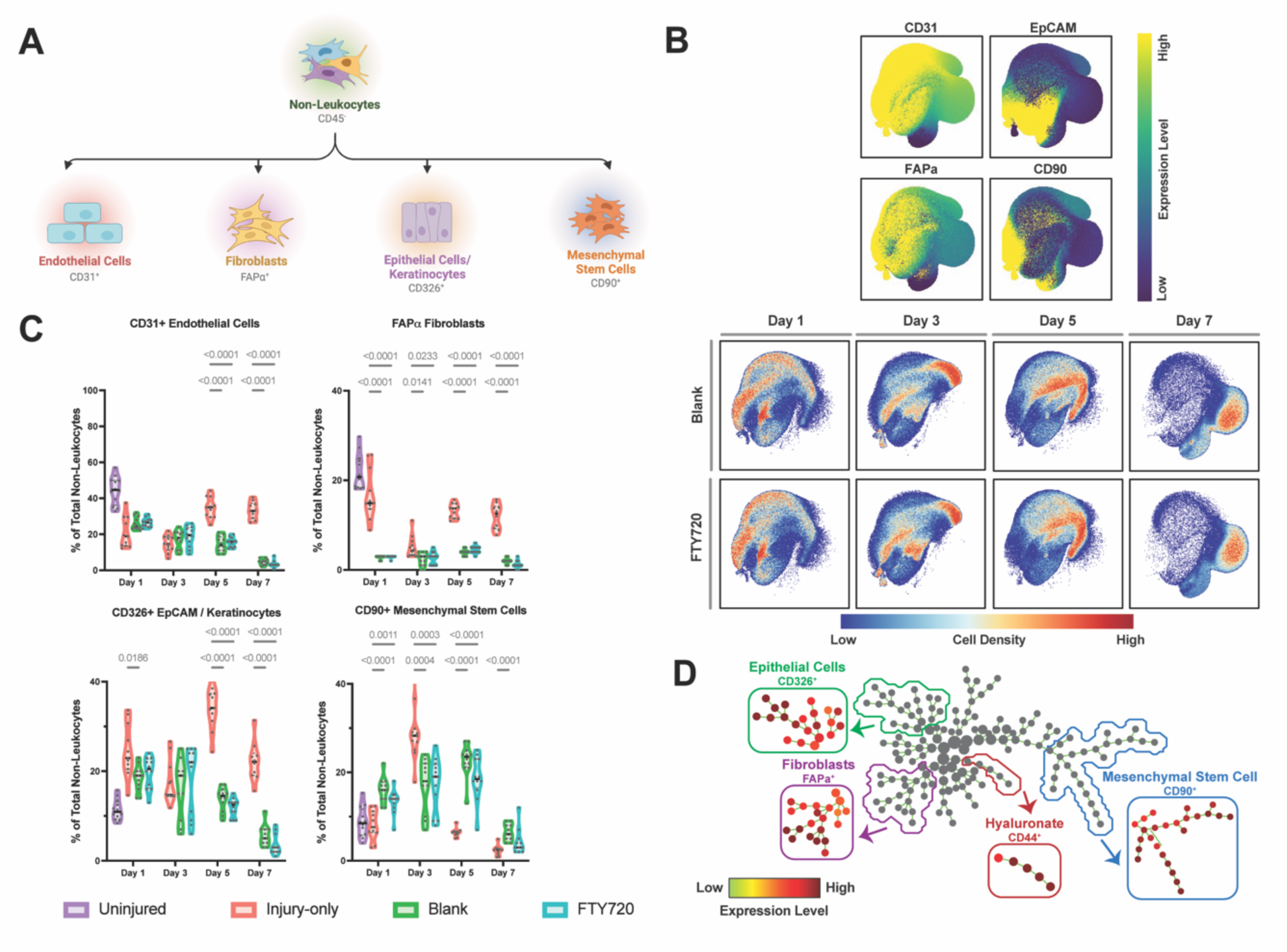
FTY720-NF reduces chronic wound response characterized by decreased stromal cell recruitment compared to injury-only mice. **A)** Gating strategy used within flow cytometric studies to identify endothelial cells, fibroblasts, keratinocytes, and mesenchymal stem cells deriving from stromal cell class. **B)** UMAPs provide a visual representation of single cells by their stromal cell expression, ranging from dark violet (low expression) to yellow (high expression) and stromal cell density following blank or FTY720-NF treatment on Days 1, 3, 5, and 7, ranging from blue (low expression) to red (high expression). **C)** Throughout all time points, there were no significant differences in stromal cell recruitment between blank and FTY720-NF groups. However, there was significantly higher FAPα^+^ fibroblast counts in injury-only mice (*P*<0.0001) and CD90^+^MSCs when compared to scaffold-receiving mice (blank, *P*<0.0001 and FTY720-NF, *P*<0.0011) on D1. Endothelial cell and keratinocytes vastly differed by D5 and D7 where the injury-only mice displayed higher percentages compared to the scaffold-treated groups (*P*<0.0001). Significance was evaluated in a 2-way ANOVA with Tukey’s multiple comparison with *P*<0.05 for n=11-12 biologically independent C57BL/6J mice per experimental group.

UMAPs were constructed using OMIQ.ai of all CD45^−^ stromal cells from blank and FTY720-NF treated mice across all timepoints. Surface marker expression values among lymphoid cell types are represented on the UMAP, ranging from dark violet (low expression) to yellow (high expression) (*Fig. 5C*). CD45^−^ non-leukocyte cellular events are represented as a dot on the UMAP with distinct cell phenotype between mice treated with FTY720-NF compared to control, ranging from blue (low cell density) to red (high cell density). However, there was no qualitative shift in stromal cell expression between mice treated with blank scaffolds compared to FTY720-NF.

Stromal cell counts from flow cytometric counts were analyzed across all timepoints for C57BL/6J mice (n=11-12) in all treatment groups: injury-only, blank, and FTY720. Cell counts were analyzed as percentage of the non-leukocyte population, presented as mean ± S.E.M, and statistical comparison made using 2-way ANOVA with Tukey’s multiple comparison revealed significance at *P*<0.05. Throughout all time points, there were no significant differences in stromal cell recruitment between blank and FTY720-NF. However, there were significant changes in cell recruitment between injury-only and scaffold-receiving groups. fibroblast percentages were significantly higher in injury-only mice on D1 compared to scaffold-receiving mice (*P* <0.0001) and remained significantly higher in all other time points. The injury-only mice revealed a significantly higher percentage of MSC on D1 compared to scaffold-receiving mice (blank, *P* <0.0001 and FTY720-NF, *P*<0.0011), but by D5 and D7, there was significantly lower MSC recruitment in the injured group (*P* <0.0001). At D5 and D7 there was significantly higher percentage of keratinocytes in injury-only mice compared to scaffold groups (blank and FTY720-NF, *P* <0.0001). Our data suggest that the nanofiber matrix alone helped regulate excessive stomal cell presence in the wound site compared to injury-only, which produced a higher percentage of fibroblasts and keratinocytes, coinciding with chronic inflammation that inhibit wound closure. Upon activation, keratinocytes express an abundance of cytokines and chemokines to transmit both positive and negative signals to cells of innate and adaptive immunity. However, it is also possible that dysregulation of keratinocytes further leads to abnormal expression of inflammatory mediators akin to pathogenesis of chronic inflammation, which further emphasizes our use of a scaffold to dampen excessive stromal cell recruitment(*33*).

A SPADE dendrogram was generated from all CD45^−^ stromal cells isolated from palate tissue of mice treated with blank or FTY720-NF at Day 1, 3, 5, and 7 (*Fig. 4E*). The stromal cell SPADE dendrogram is color depicted into four subpopulations of cells containing CD326^+^ EpCAM cells, FAPα^+^ fibroblasts, CD44^+^ hyaluronate receptor, and CD90^+^ MSC. SPADE dendrograms depict clustering of fibroblasts and epithelial cells towards the left, whereas hyaluronate receptor and MSC clustered towards the left. There remained unidentified nodes that are of CD45^−^ stomal cell lineage that may also contribute to wound healing.

### Local delivery of FTY720 sustains safe, efficacious levels for tissue biodistribution

In order to bind to S1P receptors, FTY720 is phosphorylated in vivo to its active metabolite, FTY720P, in a reaction that can be reversible or can reach equilibrium with a long elimination half-life of ∼7 days(*34, 35*). In vivo FTY720 and FTY720P concentrations were analyzed in the palate, liver, and at 2H, 6H, 1d, 2d, 3d, 5d, and 7d following bilayer scaffold implantation (*Fig. 6A*). Due to burst release and rapid initial tissue absorption of the drug upon delivery, higher concentrations of FTY720 were measured in the palatal tissue (15.52µg ± 12.93) at 2h compared to FTY720P (0.01µg ± 0.01). FTY720 concentration rapidly decreased to 0.95µg by 24hr in the palate, whereas the liver reached maximum FTY720P concentration (25.44ng ± 5.27) at 24hr with concentration decreasing to 7.08ng ± 6.306 by D7. Serum concentration of FTY720 was additionally isolated from blood and measured using LC/MS. However, FTY720 concentration was below detectable levels during LC/MS analysis, suggesting that FTY720 acted locally in the palate tissue and was not present in significant levels in the serum. A pharmacology review submitted by Novartis Pharmaceuticals established that a single oral dose of 50mg/kg FTY720 was lethal in mice, which is ∼1mg FTY720 in a 20g mouse. FTY720 was relatively well tolerated in mice up to 5mg/kg/day and the no-observed-adverse-effect level (NOAEL) was 0.5mg/kg/day(*36*). The toxicology report substantiates that our delivery of FTY720-NF was significantly below toxic levels and near NOAEL for mice. Our findings demonstrate that low dose delivery of FTY720 was sufficient to induce a wound healing response in the oral palate without detectable systemic absorption.

**Fig. 6:**
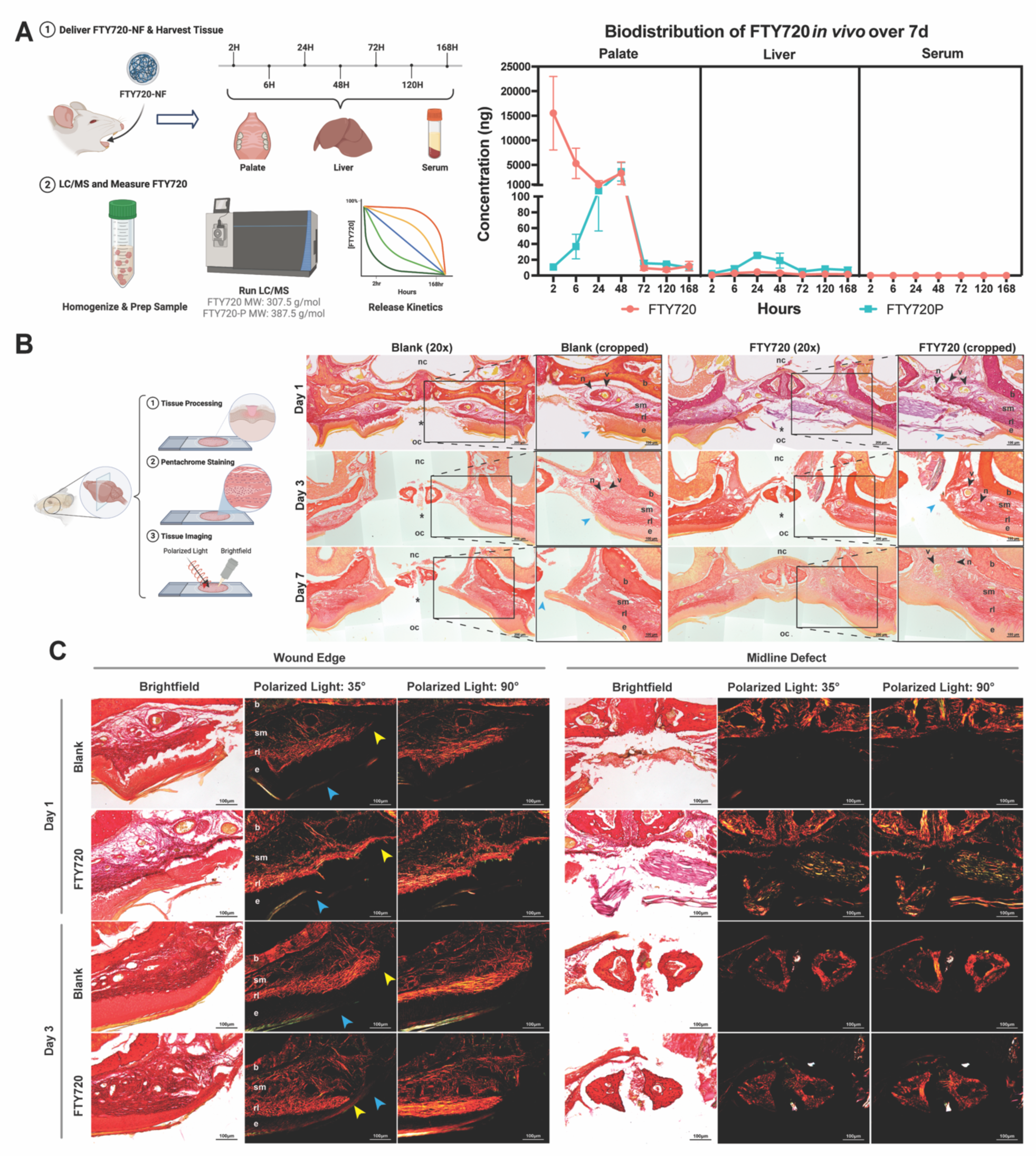
Biodistribution of FTY720 shows clearance out of palate to liver over 7d and Pentachrome staining reveals complete ONF closure following FTY720-NF treatment. **A)** Palate, liver, and blood samples were collected from C57BL/6J mice (n=3 per timepoint) treated with FTY720-NF over a 7d time course. Concentrations of FTY720 and FTY720P were analyzed at local and distal tissue via LC/MS. Peak FTY720 was observed at 2h in the palate with 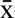 =15.52µg of FTY720 compared to 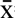 =0.01µg of FTY720P. By 24h, FTY720 concentration rapidly decreased in the palate to 0.95µg; whereas the liver reached maximum FTY720P concentration (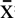 =25.44ng) at 24hr with the concentration decreasing to 7.08µg by D7. These data demonstrate that a low dose of FTY720P found in the palatal tissue by 24hr was sufficient to induce a wound healing response without detectable systemic absorption. **B)** Following Pentachrome staining to assess wound edge closure, FTY-treated mice had more medial extension of the epithelium over the exposed submucosa, forming a complete seal by Day 7; whereas blank mice had inadequate extension of the healing epithelium to reach the midline. **C)** FTY720-treated epithelium advances medially more evenly, forming a cap around the edge of the lamina propria and had more organized collagen fibers compared to control. n=3-5 biologically independent C57BL/6J mice per experimental group.

### Pentachrome staining reveals complete ONF closure and collagen alignment at wound edges following FTY720-NF treatment

FTY720-NF and blank-treated murine skull were collected at D1, D3, and D7 for histological analysis (*Fig. 6B*). Using a modified Pentachrome stain, the overall morphology and collagen fiber orientation could be resolved in the coronal plane. The oral mucosa is characteristically organized into clear layers composed of the epithelium and lamina propria (reticular layer, submucosa) which interfaces with the periosteum-lined bone. At D1, the midline defect is prominent and irregular in both treated and blank mice. The wound edge is ragged and disrupts the epithelial and submucosal layers. Both blank and FTY720-NF mice show evidence of wound healing by D3 with squamous epithelium migrating over the wound edge. However, FTY720-NF mice have more medial extension of the epithelium over the exposed submucosa, forming a complete seal by D7. Blank mice had inadequate extension of the healing epithelium to reach the midline.

Imaging of the ONF wound edge under polarized light revealed collagen fiber formation and organization (*Fig. 6C*). The collagen fibers of the disrupted oral mucosa at D1 are detached and disorganized at the medial aspect of the reticular layer, with little collagen bundling within the submucosa. By D3, collagen fibers can be seen reattaching in several orientations to the underlying bone in blank mice ahead of the squamous epithelium edge. In comparison, FTY720-NF treated epithelium advances medially more evenly, forming a cap around the edge of the lamina propria. Collagen fibers are organized into clear layers, with clear bundling and horizontal alignment within the reticular layer under 90-degree polarization.

## DISCUSSION

Pediatric patients have an unmet clinical challenge of ONF that requires repeated often unsuccessful surgeries to repair the cleft palate. Gold standard graft treatments largely act as membrane barriers and do not actively promote functional tissue regeneration or modulate the immune system towards a more pro-regenerative oral microenvironment. Current oral wound therapies have critical technological limitations including 1) the lack of an FDA-approved mucosal graft, 2) an immunomodulatory intervention via pro-regenerative recruitment of immune cells, and 3) functional barriers that lead to successful wound closure. To address these shortcomings, we engineered a Tegaderm™ bilayer, biomaterial system to provide controlled delivery of an FDA-approved, repurposed drug, FTY720, to effectively modulate wound healing towards a pro-regenerative phenotype following ONF injury. This bilayer system sustains local release of FTY720 with safe tissue biodistribution, significantly enhances ONF healing and wound closure, and modulates the immune response to drive tissue regeneration. By identifying the optimized dosing level of 1:60 drug:polymer ratio, we were able to design the FTY720-NF to sustain release of drug for up to 7d in vitro. In developing a bilayer system using occlusive Tegaderm™ wound dressing, we successfully implanted the scaffold in vivo to the ONF injury and maintained local, 8–12-hour transdermal drug release that allowed for complete wound healing by D7. We provided extensive characterization of local oral immune responses to FTY720-NF and identified a state transition in both myeloid and lymphoid immune cell subsets from inflammatory to pro-regenerative phenotypes that facilitated wound closure via stromal cell recruitment (*Fig. 7*).

**Fig. 7:**
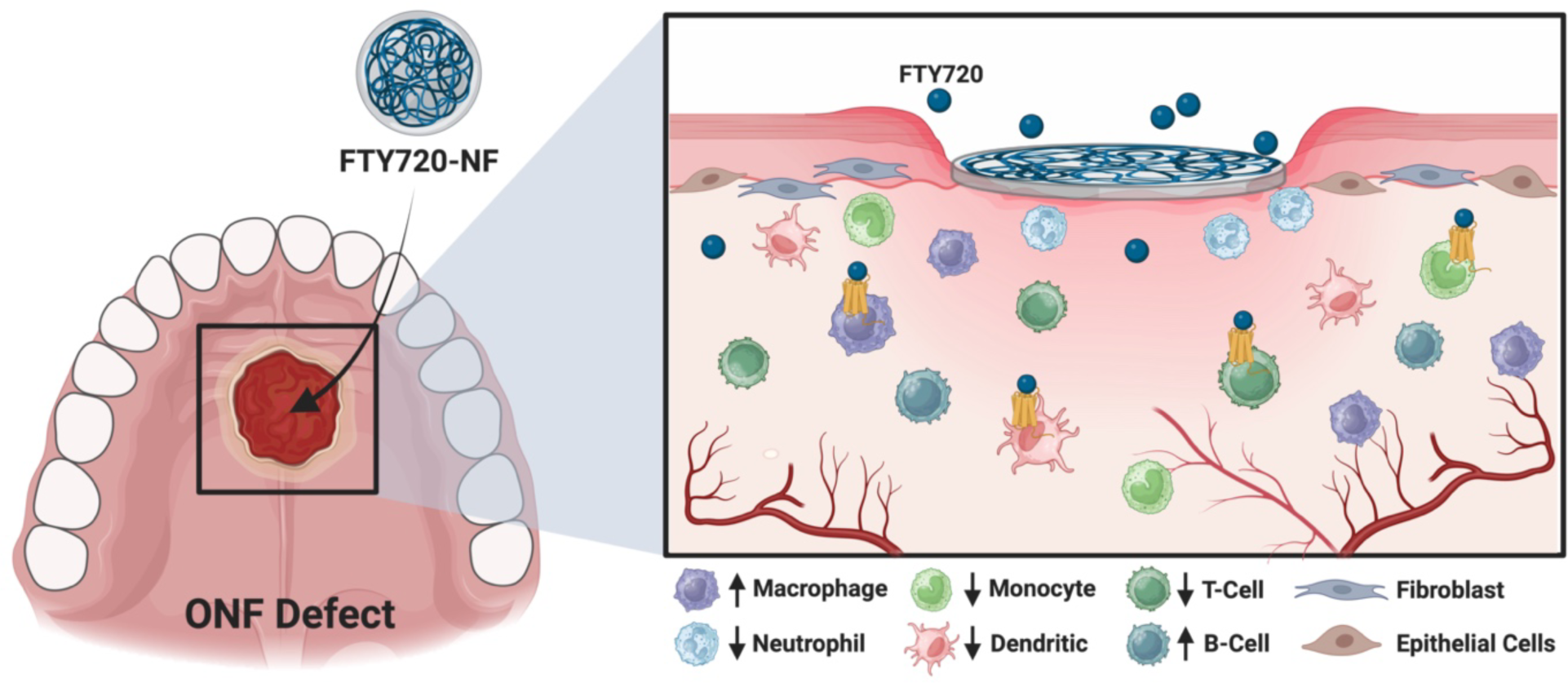
FTY720-NF was implanted in vivo to mice with ONF injury on the hard palate mucosa. Local delivery of FTY720 modulated S1P receptors to decrease recruitment of neutrophils and monocytes at D1 and increase macrophage recruitment on Day 3. Treatment with FTY720 revealed a decrease of lymphoid-derived DCs on D1 and D3. There was a decrease in T cell recruitment on D5 with increases in B cell percentage on D7. Treatment with nanofiber scaffold downregulated aberrant stromal cell infiltration to avoid overproduction of fibroblasts and epithelial cells that would be detrimental to wound closure.

Our comprehensive immunophenotyping data support what is known about the oral wound healing cascade and provide greater understanding of the function of FTY720 to help resolve oral wounds following an ONF injury. FTY720 caused significantly less neutrophil and monocyte infiltration in the palate on D1 following injury, likely because FTY720 restricts neutrophil and monocyte egress from the bone marrow via S1PR_1/4_ and S1PR_3,_ respectively(*37, 38*). Our neutrophil and monocyte SPADE dendrograms provide new insight into the pseudotime shift in immune cell state, which illustrate that treatment with FTY720 transitioned Ly6G^hi^ neutrophils towards a hybrid CD62L^+/−^ state and Ly6C monocytes towards a Ly6C^lo^ anti-inflammatory phenotype by D7. In contrast, FTY720-NF treated mice presented an influx of MerTK^+^ macrophages on D3, but no significant difference in the polarization of macrophages. Macrophages are widely known to facilitate the maintenance and resolution of inflammation by downregulating inflammatory factors and upregulating anti-inflammatory growth factors that send out activation signals to lymphoid and stromal cells(*39*). Previous studies have shown that S1P conventionally promotes CD86^+^ M1 macrophage polarization through SIPR_3_, but as an S1P-antagonist, FTY720 attenuates CD206^+^ M2 macrophage polarization instead(*37*). Although we did not see a significant difference in FTY720-mediated macrophage polarization in our flow cytometric analyses, our SPADE dendrogram illustrate that by D7, macrophages indeed polarized to M1, M2, or hybrid M1/M2 phenotypes. It is possible that the hybrid macrophage phenotype helped to resolve inflammation which in turn helped to drive ONF repair, similar to a study where S1PR modulators induced hybrid macrophage polarization in a murine volumetric muscle loss model(*18*). In follow-up studies, we can continue to explore the role by which FTY720 modulates macrophages in the context of ONF healing and modifies their role toward a pro-regenerative phenotype.

Our quantitative data suggests that treatment with FTY720 does not affect all lymphocytes equally. We provide evidence from DC SPADE dendrograms that suggest that the lineage from which DCs originate dictates the functions by which ONF healing occurs. Prior studies have shown that classical DCs can be lymphoid CD11b^−^ CD8^+^ or myeloid CD11b^+^ derived and such sublineage allow DCs to form a unique transcriptional entity(*40*). CD8^+^ DCs are known to direct the development of T_h_ responses and are specialized for phagocytic uptake of dead cells. Conversely, CD11b^+^ DC differentiation is controlled by granulocyte-macrophage colony-stimulating factor (GM-CSF) and are specialized for presentation on major histocompatibility complex class II(*41*). Our dendrograms show a clear upregulation in lymphoid derived DCs, indicating that ONF healing required differentiation of CD8^+^ DCs. These lymphoid derived DC are stimulated by cytokine FMS-related tyrosine kinase 3 ligand (FL), rather than GM-CSF, which has major influences on the development of inflammatory DCs(*42*). Furthermore, we saw significant decreases in the percentage of CD3^+^ T and CD4^+^ T_h_ cells at D5 and D7, which is in line with other studies showing that FTY720 dramatically suppresses T-cell egress from the lymph node via S1PR_1_ to prevent hyper-activation of lymphoid cells contributing to a pro-inflammatory response(*43, 44*). However, it was striking that B cell percentages were increased following treatment with FTY720 considering that B cells are also modulated by S1PR_1_. Similar diminished effects of FTY720 on B cells were seen in a mouse model of MS, suggesting that the increased B cell production could be due to 1) less dependence on S1P_1_-mediated egress compared to T cells and 2) their relative resistance to apoptosis induction by FTY720(*45, 46*).

Our previously published study showed that FTY720-NF promotes oral wound closure but did not explore the extent of stromal cell function in tissue regeneration(*14*). Unlike previous studies that developed scaffold delivery systems for pre-clinical oral wound models, we were able to provide both quantitative and qualitative insights in stromal cell functions in the context of nanofiber scaffolds for treatment of ONF healing(*2, 47–50*). The crosstalk between fibroblasts and leukocytes forms a powerful cell-intrinsic amplification loop that can give rise to pathological overproduction of fibroblasts, which have been found to be central drivers of fibrotic conditions in nonhealing wounds due to the release of inflammatory cytokines and chemokines(*51*). Such autocrine amplification loop of fibroblasts can lead to pathological behavior stemming from overproduction of pro-inflammatory factors often seen in chronic wounds as reported in chronic diabetic foot ulcers(*52*). We highlight that by incorporating a nanofiber scaffold at the wound site, we can downregulate aberrant stromal cell infiltration to avoid overproduction of FAPα^+^ fibroblasts and CD326^+^ EpCAM/keratinocytes that would be detrimental to wound closure evidenced in injury-only mice compared to scaffold-receiving mice. Exclusive to our pentachrome histology, we showed robust, full-thickness wound closure by D7 following TD bilayer delivery of FTY720-NF when compared to control groups. Furthermore, we were able to identify that collagen synthesis plays a pivotal role in dictating quality of tissue regeneration, structural integrity of healed tissue, and matrix remodeling. To understand the role of collagen fibers in wound closure, we utilized polarized light microscopy to show that mice treated with FTY720-NF had epithelium advance medially and evenly around the edge of the lamina propria with organized collagen fibers compared to control. These findings provide critical translational support for the further development of immunomodulatory therapeutics for oral wound healing, potentially offering greater opportunities for an FDA-approved treatment option for mucosal tissue repair.

## MATERIALS AND METHODS

### FTY720 Scaffold Nanofiber Fabrication and Electrospinning

Electrospun nanofiber scaffolds were fabricated with or without FTY720 drug solution for dosing study and all subsequent in vivo scaffold implantation studies. Polymer blend of 1:1 weight ratio of 75:25 poly(lactide-co-glycolide acid) (Sigma Aldrich #719927) and polycaprolactone (Sigma Aldrich #440744) was dissolved in glass scintillation vial with 2ml solution of 1:3 Methanol: Chloroform solvent. FTY720 (US Pharmacopeia) drug at 5mg/ml and 10mg/ml concentration were dissolved in Methanol: Chloroform solvent and thoroughly mixed with polymer blend solution under constant sonication and shaking for 2h. Polymer+drug solution (2mL) was electrospun 1ml/hr rate at 19kV using a Harvard Apparatus PHD 2000 infusion pump and HV Power Supply (Gamma High Voltage Research #ES30P-5W/DAM). Nanofiber sheets were lipolyzed overnight and stored at −80°C until use.

### In Vivo ONF Injury Murine Model and Scaffold Implantation

Male C57BL/6J mice (Jackson Laboratory #000664) of age 4-6 weeks were used for all studies. Mice were anesthetized with a Ketamine (100mg/kg): Xylazine (10mg/kg) cocktail intraperitoneally, and Buprenorphine SR (0.5mg/kg) was administered subcutaneously as an analgesic. ONF injury of 1.5mm full thickness defect was modeled as a critically sized defect between the 3^rd^ and 4^th^ rugae in the hard palate mucosa of mice using a High Temperature Cauterizer (Bovie Medical Corporation). A 1.5mm biopsy punch was used to ensure consistent size of the injury between the mice. Blank and FTY720-lNF scaffolds were implanted at sites immediately after ONF injury and secured using a tissue adhesive. ONF healing was monitored and imaged Days 1, 3, 5, and 7 post-ONF formation using an Endoscope (SKOR). On the day of tissue harvest, mice were euthanized via CO_2_ asphyxiation and cervical dislocation. Hard palate mucosa was excised using a scalpel with #11 straight, angled blade (Integra Miltex #4-111) and blood was collected in K2 EDTA-coated tubes (BD #367841) via cardiac puncture and samples were stored appropriately for further analysis. Sample size was determined using a power analysis from our pilot study, indicating at least 6 samples per treatment group. All surgical procedures were performed in compliance with Emory University’s Department of Animal Research and Institutional Care and Use Committee.

### FTY720-NF Release Kinetics and Drug Loading Efficiency

Biopsy punch was used to create 3mm diameter nanofiber scaffolds. FTY720-loaded NF were placed in simulated body fluid containing 5% fatty acid-free bovine serum albumin in phosphate buffered saline (PBS). Releasate was collected and replaced with new stimulated body fluid at 0hr, 1hr, 2hr, 6hr, 24hr, 48hr, 3d, 5d, and 7d timepoints. The remaining scaffold was dissolved in 1:3 Methanol:Chloroform solution to obtain total drug loaded into each scaffold. Equal volumes of 100% Acetonitrile (Sigma-Aldrich #34851) were added to releasate and relative abundance of FTY720 was measured using LC/MS courtesy of Dr. Dennis Liotta’s Mass Spectroscopy facilities at Emory University Department of Chemistry.

### Scanning Electron Microscopy (SEM)

Electrospun nanofibers loaded with/or without FTY720 were cut into 1.5mm diameters scaffolds using a biopsy punch. Scaffolds were sputter coated with gold for 30s and SEM images of blank, 5mg, and 10mg scaffolds were taken at 300x, 1000x, and 3000x zoom at 15kV using Emory University’s *Robert P. Apkarian Integrated Electron Microscopy Core*.

### Atomic Force Microscopy (AFM)

All atomic force microscopy (AFM) measurements were performed by Dr. Alan Lui curtesy of Dr. Todd Sulchek Lab at Georgia Tech’s Department of Mechanical Engineering. Electrospun nanofibers loaded with/or without FTY720 were cut into 8mm diameters scaffolds using a biopsy punch. Nanofibers were fixed on glass slides with double-sided tape. For stiffness measurement, 5.24um um Si beads were attached to tipless silicon cantilevers using two-part epoxy and dried for at least 24hr. The thermal method was used to calibrate the cantilever spring constant immediately prior to use by indenting a glass substrate and performing a Lorentzian fit to the thermal spectrum. Topological images and stiffness characterizations were obtained from 80 × 80 μm^2^ force maps with 96 × 96 force curves per image. All the force-distance measurements were performed with 2 μm/s approach velocity using a 500nN force trigger. Force-distance plots were transformed into force-indentation depth plots, then 10–100% indentation depth was used to evaluate Young’s modulus using a Hertzian contact model.

### Endoscope Images and Wound Closure Analysis

Following ONF injury, palatal images are taken by an endoscope (SKOR) at Level 3 brightness on Days 1,3,5, and 7. The mice’s jaw is propped open an eyelid retractor and a camera probe was held approximately 4 inches above the palate for full hard palate view. The oral palate was measured to be approximately 4.5mm in length between the side molars. Images and injury sizes were analyzed using ImageJ. Percent Healed Area was measured using the following formula: 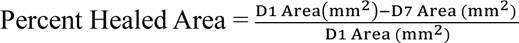.

### Scaffold Adhesive Methods

Various adhesives were purchased to secure and test the efficacy of FTY720-NF adhesion to the hard palate mucosa. Following ONF injury in the hard palate as described previously, NF scaffolds were inserted at the site of injury and one of the following adhesives was utilized to attach the scaffold to the surrounding oral mucosa. Endoscope images were taken of the scaffold and adhesive on the palate and the time of scaffold dissociation was recorded.

### Palate Tissue Staining and Flow Cytometry

All flow cytometric studies and subsequent analyses were conducted with n=11-12 biologically independent C57BL/6J mice per experimental group (*Supplemental Fig. 2A*). Fresh palatal mucosa was harvested, and tissue was processed in digest solution containing Collagenase II 5500U/mL (Thermofisher #17101015) and Dispase II 2.5U/mL (Sigma #D4693-1G) for 60-80min in 37°C water bath. Single cells were washed in ice cold PBS and centrifuged at 400g for 5min with solution discarded afterward. Similarly, blood collection from each mouse was processed using red blood cell lysis buffer and filtered to obtain a white cell pellet of leukocytes. Both palate and blood cells were stained with viability staining, and cells were incubated for 15min in mouse Fc block solution (BD Biosciences #101301). Cells were stained for 40min in master mix antibody cocktail using antibody concentrations optimized for this flow panel (*Supplemental Fig. 2B*). Cells were fixed in 4% paraformaldehyde for 15min, washed in FACS buffer, and stored overnight in 4°C until flow cytometry run. All flow runs were carried out by 5-Laser Cytek Aurora curtesy of Emory Pediatrics/Winship Flow Cytometry core and single cells were selected in OMIQ.ai for all subsequent analysis.

### Uniform Manifold Approximation and Projection (UMAP)

UMAP was performed using OMIQ.ai software as a nonlinear dimensionality reduction algorithm. UMAP can embed high-dimensional data into a space of two or three dimensions, and cells are visualized in a scatter plot, where similarity is determined via proximity to other points. In preparation for UMAP, each flow cytometry sample was pre-gated to select immune cell class (i.e., myeloid, lymphoid, stromal, neutrophils) and then imported into OMIQ.ai. UMAP parameters of n_neighbor= 15, min_distance= 0.4 were applied. UMAP projection was generated for cellular markers of interest, and samples from specific treatment groups or timepoints were visualized by overlaying scatterplots.

### Spanning-tree Progression Analysis of Density-normalized Events (SPADE)

SPADE was performed through MATLAB and the source code is available at http://pengqiu.gatech.edu/software/SPADE/. SPADE is a visualization tool creating a 2D minimum spanning tree organized into nodes representing clusters of cells similar in their surface marker expressions. The size and color of each node are relative to the number of cells present and the median marker expression. SPADE automatically generates the tree by performing density-dependent downsampling, agglomerative clustering, linking clusters with a minimum spanning-tree algorithm and upsampling based on user input. The SPADE tree was generated by exporting compensated select cellular subsets (i.e., CD3^+^ T cells). The following SPADE parameters were used: ignore compensation, no transformation, neighborhood size 5, local density approximation factor 1.5, max allowable cells in pooled downsampled data 50000, target density 20000 cells remaining, and number of desired clusters of 100.

### Histology

Mice heads were skinned, fixed in 4% paraformaldehyde for 48h, and decalcified with Cal-Ex decalcifier (Fisher Scientific #CS510) for 5d. Following decalcification, samples were dehydrated, infiltrated with Paraplast paraffin wax (Leica, #39601006), and embedded for coronal sectioning. 7-10µm sections were mounted on charged glass slides for further histochemical staining with a modified pentachrome method. Briefly, sections were deparaffinized and rehydrated through graded ethanol washes to water, followed by pre-treatment with 6% nitric acid and sequential staining with 0.1% toluidine blue (EMS #26074-15) and picrosirius red (Abcam, #ab246832). Slides were then dehydrated and cover-slipped with toluene-based mounting medium for imaging.

### Microscopy

Brightfield images were captured with an AxioScan.Z1 slide scanner (Zeiss) equipped with a Plan-apochromat 40X/0.95 objective. Darkfield polarized transmission images were captured with a 10X/0.3NA objective using an Olympus BX51 microscope (Olympus) equipped with a condenser with NA adjusted to 0.3, a rotatable sample stage, and an Infinity 2C color camera (Luminera). The rotatable polarizer and polarizer-analyzer were crossed at 90° through the microscope light path.

### Biodistribution

Palate, liver, and blood samples were collected from mice with FTY720 (n=3) at 2H, 6H, 1d, 2d, 3d, 5d, and 7d. 50µL of serum was mixed with 100µL of ice-cold acetonitrile containing internal standard (d5-ethoxycoumarin), vortexed for 10 seconds and centrifuged at 13,000RPM at 4°C for 10mins to pellet proteins. Supernatants were transferred to LC-MS autosampler vials for instrumental analysis. Tissue samples were extracted with ice-cold acetonitrile containing internal standard (d5-ethoxycoumarin) based on wet tissue weights. Liver samples were extracted with the addition of 9µL of acetonitrile per mg of tissue and palate samples were extracted with the addition of 24µL of acetonitrile per mg of tissue. Tissue samples were homogenized with ceramic beads (OMNI Biosciences) added to each sample using a Lysera homogenizer (Biotage) set at 4°C. Tissue homogenates were centrifuged at 13,000RPM at 4°C for 20 minutes to pellet proteins and other insoluble fractions. Supernatants were transferred to autosampler vials for instrumental analysis. Calibration curves were prepared with the addition of known concentrations of FTY720 and FTY720P from matrix-matched control serum, palate, or liver samples. Samples were analyzed with LC-MS/MS analysis using an Agilent 1260 LC coupled to a 6460C Triple Quadrupole mass spectrometer.

### Sample Size and Statistical Analysis

All statistical analysis was performed in GraphPad Prism software. Data presented as mean ± S.E.M. unless otherwise indicated. *A priori* power analysis was run to determine appropriate study sample size for mice receiving no scaffold (-CTRL), blank scaffold (+CTRL), and FTY720-NF (treatment). Power analysis was run with power=0.8 and α=0.05, using wound size as the primary outcome measure for 7d mice studies. For comparison of two groups, an unpaired *t* test was used. For comparison between oral wound size across time, Kruskal-Wallis one-way ANOVA was used. For grouped analyses, significance difference between was determined using two-way ANOVA with Tukey post-hoc test for multiple comparison. Correlation analysis, such as multivariate dimensionality-reduction tools were used to determine significant relationships between in vivo data sets. Statistical significance is set at *P*<0.05.

## ACKNOWLEDGEMENTS

AIT designed performed the experiments, analyzed the data, and wrote and revised manuscript. AIT and DS were responsible for analyzing flow cytometry data and performing dimensionality reduction analyses. DR, JOP were responsible for processing animal samples for pentachromic histology and microscopy imaging. LH, TT, AK assisted in protocol training and experimental designs. KL, PB, LJ performed MS/LC for drug dosing and biodistribution studies. Figures were created using BioRender.com and Adobe Illustrator. Revising and approving final version of manuscript: All authors. All authors have read the journal’s authorship agreement and declare no conflict of interest. Research reported in this publication was supported by the Oral Maxillofacial Surgery Foundation (Funding ID: 2591) and the National Institute of Dental and Craniofacial Research of the National Institutes of Health under award number R01DE028905. The content of this article is solely the responsibility of the authors and does not necessarily represent the official views of the National Institutes of Health. This manuscript has not previously been copyrighted or published and is not under consideration for publication elsewhere.

