## Supplemental 1 for "Harnessing Bilayer Biomaterial Delivery of FTY720 as an Immunotherapy to Accelerate Oral Wound Healing"

**A**

| Potential Adhesive Methods | | |
|---|---|---|
| **Product Name** | **Manufacturer** | **Key Ingredient** |
| Vetbond | 3M | Cyanoacrylate |
| LiquiVet Tissue Adhesive, 8 Seconds | Oasis | Cyanoacrylate |
| Ora-Aid | TBM Corporation | Hydroxyl ethyl cellulose |
| Tegaderm Transparent Dressing TEGA-10 | Nexcare | Polyurethane, acrylic adhesive |
| Band-Aid Hydro Seal Hydrocolloid Bandage | Johnson & Johnson | cross-linked matrix gelatin, pectin, and carboxymethyl-cellulose |
| Super Poligrip Comfort Seal Denture Adhesive Strip | GlaxoSmithKline | PEG-90M |
| 8P-T Mucosal Hydrogel | Messersmith Research Group, UC Berkley | Tren terminated 8-arm-PEG-10k |

**B**

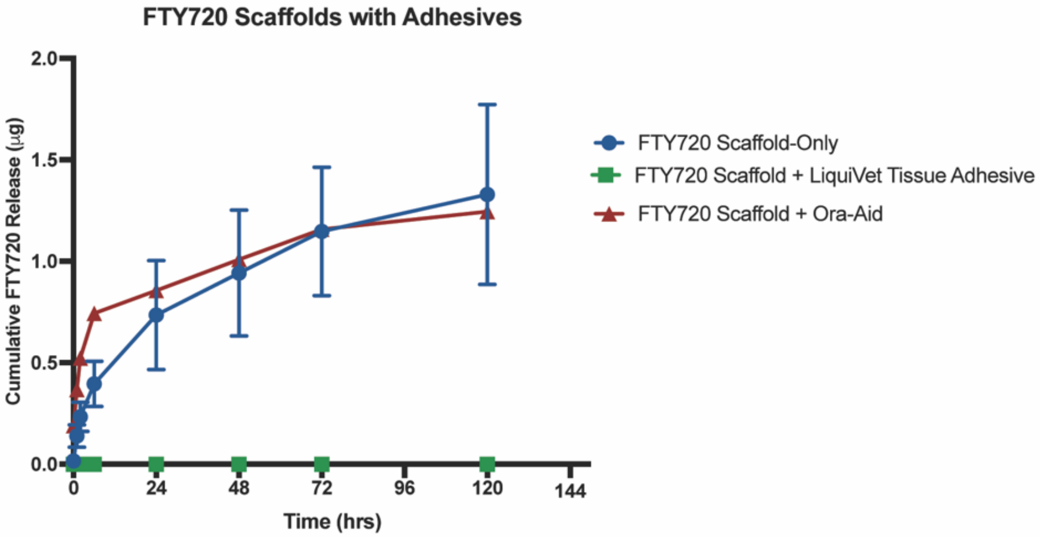


**Supplemental 1: A)** List of potential Adhesive methods used to help retain scaffold attachment to oral mucosa. Adhesion methods ranged from wound dressing to polymer-based materials. **B)** LC/MS measures of cumulative drug released following attachment to scaffold *in vitro*. Scaffold directly attached using LiquiVet tissue adhesive glue did not release drug over 5d time course.

**Title: Harnessing Bilayer Biomaterial Delivery of FTY720 as an Immunotherapy to Accelerate Oral Wound Healing**

**A**

| Treatment Groups | Day 1 | Day 3 | Day 5 | Day 7 |
|---|---|---|---|---|
| Uninjured | 12 | | | |
| Injury-Only | 12 | 12 | 12 | 12 |
| Blank-NF | 12 | 12 | 12 | 12 |
| FTY720-NF | 12 | 12 | 12 | 12 |

**B**

| Emory Flow Cytometry Core: Cytek Aurora Flow Panel | | | |
|---|---|---|---|
| **Leukocytes** | CD45-PerCP Cy5.5 | **B-Cells** | CD19-BUV615 |
| | CD11b-BUV563 | **T-Cells** | CD3- Pacific Blue |
| **Dendritic Cells** | CD11c- BUV805 | **Helper T-Cells** | CD4-BV570 |
| **Monocyte & Macrophages** | CD64-PE-Cy7 | **Cytotoxic T-Cells** | CD8-AF700 |
| | Ly6C-BV421 | **Fibroblast** | FAPa- AF488 |
| | MerTK-BB700 | **Keratinocytes** | EpCAM/CD326- BV711 |
| | CD206-PE Dazzle 594 | **Vascular Endothelial Growth Factor** | CD31-BV510 |
| | CD86-BV650 | **MSC Cells** | CD90-BV786 |
| **Neutrophils** | Ly6G-APC | **HSC/Epithelial Progenitor** | CD34- AF532 |
| | CD62L-BUV737 | **Hyaluronate Receptor** | CD44-BV605 |
| **Natural Killer Cells** | CD161-BUV496 | **Viability** | Zombie NIR-APC/Cy7 |

**Supplemental 2: A)** Sample size of n=12 biologically independent C57BL/6J mice we used per experimental group for flow cytometric studies. **B)** List of flow cytometry markers and fluorophores that was optimized for Aurora Cytek 5L at Emory Pediatrics/Winship Flow Cytometry core.
